# DNFE: Directed-network flow entropy for detecting the tipping points during biological processes

**DOI:** 10.1101/2024.09.18.613673

**Authors:** Xueqing Peng, Peiluan Li, Chen Luonan

## Abstract

There generally exists a critical state or tipping point from a stable state to another in dynamic biological processes, beyond which a significant qualitative transition occurs. Identifying this tipping point and its driving network is essential to prevent or delay catastrophic consequences. However, most traditional approaches based on undirected networks still suffer from the problem of the robustness and effectiveness when applied to high-dimensional small sample data, especially for single-cell data. To address this challenge, we developed a directed-network flow entropy (DNFE) method which can transform measured omics data into a directed network. This method is applicable to both single-cell RNA-sequencing (scRNA-seq) and bulk data. By applying this method to five real datasets, including three single-cell datasets and two bulk tumor datasets, the method can not only successfully detect the critical states as well as their dynamic network biomarkers, but also help explore regulatory relationships between genes. Numerical simulation indicates that the DNFE method is robust and superior to existing methods. Furthermore, DNFE has predicted active transcription factors (TFs), and further identified ‘dark genes’, which are usually overlooked by traditional methods.

## INTRODUCTION

Many complex systems undergo a critical transition, from one state abruptly switching to a contrasting state^1^. This is known as a tipping point, at which a drastic, irreversible, and qualitative transition may occur. Single-cell RNA sequencing has revolutionized the study of cellular heterogeneity and functional diversity, enabling gene expression measurement within individual cells and allowing inference of cell population arrangement based on trajectory topology and gene regulatory analysis^2,3,4^. There is a tipping point during many biological processes, such as a cell differentiation process or embryonic development, after which there is a drastic transition or change in cell populations^5^. Identifying such a tipping point just before the critical transition in biological systems is crucial in providing insights into the underlying mechanisms of disease progression or embryonic development^6,7^. However, due to the similarity between the pre-transition and critical states in terms of both phenotype and mean gene expression, traditional biomarkers often fail to detect the critical state.

To signal the critical state before the transition of biological systems, a theoretical concept known as the dynamic network biomarker (DNB) was proposed8. DNBs are composed of a group of molecules, such as genes or proteins, which are strongly correlated with each other, i.e. a phenomenon called “critical collective fluctuation”. Based on the DNB theory, numerous methods have been developed and successfully applied to detect the critical states of complex diseases and biological processes, such as influenza^9^, breast cancer^10^, bladder urothelial carcinoma^11^, and cell differentiation^12,13^. However, the application in scRNA-seq data analysis is limited due to severe interference from transcript amplification noises and dropout events. Moreover, most of these computational approaches merely identify correlations among molecules/genes based on undirected networks, yet directed networks can reflect the interactions among genes, mine the underlying dynamic information in cell populations, and reveal the basic mechanism of molecular effects to provide quantitative studies for further precise treatment. A directed network can also explicitly represent directional relationships between genes, which is crucial in describing causal relationships, gaining a better understanding of the regulatory relationships between genes, and identifying important regulatory factors and pathways. Therefore, it is imperative to develop an effective and robust method for detecting tipping points based on directed networks, especially for single-cell data of a complex biological process.

To address these issues, we propose a novel computational method based on directed networks, called directed network flow entropy (DNFE), to predict the critical transition for bulk and single-cell data. Unlike traditional methods, we focus on the first-order neighbors that connect directly to the core gene, as well as the second-order neighbors that have direct interaction with any of the first-order neighbors. Specifically, we first construct a time-specific directed network at a given time point based on rewiring the WGCNA network with a direction determination index. We then calculate the local DNFE for each local directed network according to the information of the directed network. Unlike traditional information entropy, DNFE fully utilizes local (core gene) network connectivity information rather than fluctuated gene expressions, thereby reliably quantifying network fluctuation. Finally, we use the DNFE score to characterize the molecular collective fluctuation or network fluctuation caused by specific samples against the reference samples/cells, and qualify the criticality of the biological process, i.e. critical collective fluctuation. The proposed DNFE method serves as an effective tool to detect the critical state during a complex biological process, with the following advantages: (i) The DNFE method reliably quantifies critical network fluctuation, reducing noise by exploring dynamic and high-dimensional information of omics data, thereby enhancing the robustness of the method. Moreover, we focus on the second-order neighbors which better depicts the structure of the network, thereby improving the effectiveness of the method. The numerical simulation shows the algorithm’s robustness, effectiveness and applicability to handle vast amounts of data or large-scale datasets. (ii) Unlike most traditional methods, the DNFE method is proposed based on directed networks, which can help us gain a better understanding of the regulatory relationships between genes, identify important regulatory factors and pathways, and explore the structure and functionality of a gene regulatory network. (iii) The DNFE method can detect critical state before the state transitions and identify effective DNB members with critical collective fluctuations and non-differential ‘dark genes’, which may be used as prognostic biomarkers of complex diseases and biological processes. (iv) The DNFE method can be applied to both bulk and single-cell RNA-seq data. The method can also identify transcription factors (TFs), which are key players that define cell identity and drive cell-fate transitions. One of the identified such TFs is ZNF888, which was considered to hold great potential in early cell differentiation.

To demonstrate the robustness and effectiveness of the DNFE, we performed numerical simulations based on the gene expression data generated by the artificial gene regulatory network under different noise strengths. As noise strength increased, DNFE performed better in detecting early-warning signals for the upcoming tipping point compared with the existing method^14^. Furthermore, the numerical simulation, conducted on a 1000-node network, unequivocally showed the algorithm’s robustness and applicability to handle vast amounts of data or large-scale datasets. The DNFE method was applied to two bulk sequencing tumor datasets, including kidney renal papillary cell carcinoma (KIRP) and bladder cancer (BLCA) from the Cancer Genome Atlas (TCGA) database, successfully detecting early-warning signals of critical states or tipping points. Additionally, cell fate commitment was successfully detected in three single-cell datasets of embryonic development, including the differentiation of human embryonic stem cells to definitive endoderm cells, mouse ESC (mESC) to mesoderm progenitor (MP), and mouse embryonic fibroblast (MEF) to neuron. DNFE identified the two tipping points during the differentiation process based on DECs data, i.e., the first tipping point at 12h and the second tipping point at 36h, which correspond to two different differentiation processes, i.e., hESC (ES)-to-ME and ME-to-DE.

## RESULTS

### The overview of the DNFE algorithm

Given a collection of control cells/samples and a corresponding set of case cells/samples, a specific directed network is constructed. This construction is based on the rewiring of a weighted gene coexpression network analysis, guided by a direction determination index ω_ij_(Fig 1A). Subsequently, the global Directed Network Function Efficiency (DNFE) is calculated. This measure is employed to detect early-warning signals for the critical transition of a complex biological process(Fig 1B). During the dynamic progression of this biological process, the DNFE score remains low when the system is in a stable state. However, it experiences a significant increase when the system approaches the critical state. This abrupt increase in the DNFE score serves as an indicator of the tipping point in the biological process(Fig 1C). The theoretical background can be seen in Supplementary material: A.

**Figure 1.**
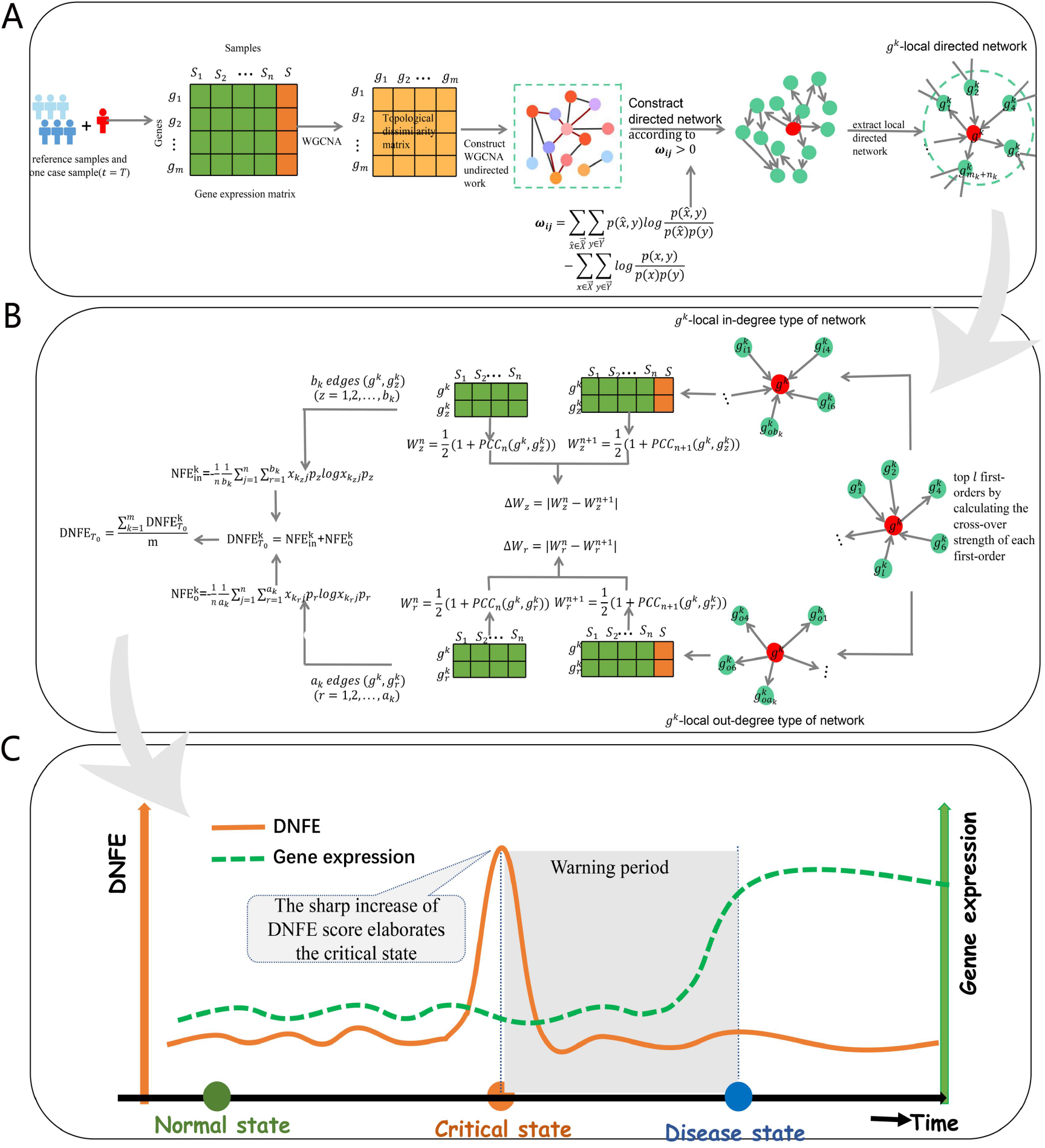
The schematic of the DNFE algorithm. (A) Given a group of control cells/samples and a set of case cells/samples derived, the specific directed network is constructed based on rewiring weighted gene coexpression network analysis with direction determination index. (B)The global DNFE is calculated, which is utilized to detect the early-warning signal for the critical transition of a complex biological process. (C)During the dynamic progression of biological process, the DNFE score remains low when the system is in a normal state, while it increases significantly when the system is close to the critical state. Such an abrupt increase in the DNFE indicates the tipping point of biological process.

### Validation based on numerical simulation

To validate the effectiveness and robustness of the proposed DNFE method, we used an eleven-node artificial network (Fig 2A) to illustrate the identification of early-warning signals as the system approaches a tipping point. Models employing Michaelis - Menten kinetics or the Hill form are typically used to represent the dynamics of gene regulatory networks^15,16,17^, such as transcription and translation^18,19^, cyclic reactions^20^, nonlinear biological processes^21,22^, and other gene regulatory activities^23,24^. Based on a parameter varying from -0.5 to 0.23 with a bifurcation parameter value as the tipping point, two simulated datasets were generated.

**Figure 2.**
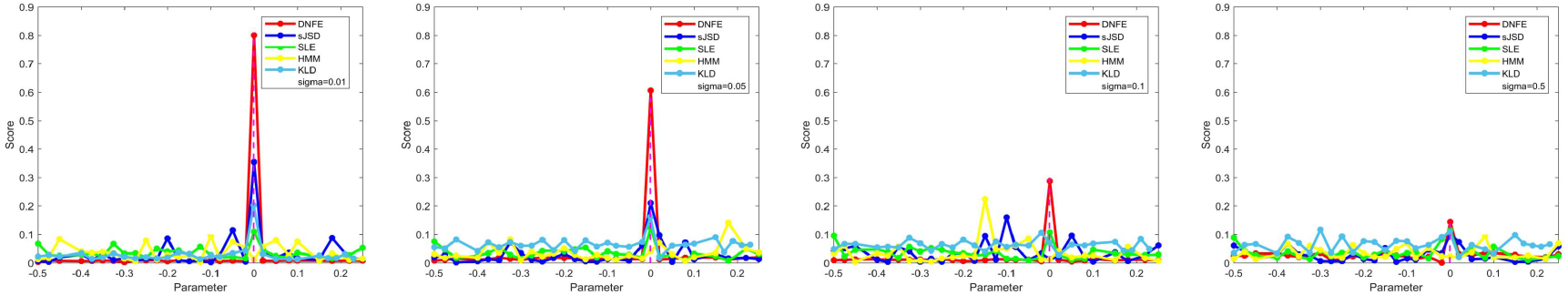
The validation of the DNFE method on a simulation dataset. (A) The model of an 11-node network, in which the arrow represents positive regulation whereas the blunt line denotes negative regulation. (B) The curve of the DNFE score based on the gene regulatory network. (C) The landscape of the DNFE scores for 11 nodes. (D) Comparison of the robustness between the DNFE method and sJSD method at different noise strength. (E) The curve of the DNFE score based on the 100-node gene regulatory network. (F) The curve of the DNFE score based on the 1000-node gene regulatory network.

Fig 2B illustrates the dynamic change in the DNFE score for the 11-node network. There is a sharp increase in the DNFE score at the parameter value, indicating the imminent tipping point or critical transition at a bifurcation point. To better exhibit the distinct dynamics of the system between the normal state and the pre-disease state, the dynamical changes were presented in local DNFE scores respectively for 11 local networks, illustrating the landscape of the network flow entropy in a global view in Fig 2C. Clearly, when the system is far from the tipping point, the DNFE scores of all nodes are steady and at a low level; when the system approaches the tipping point, some nodes behave in a considerably different manner in terms of their expression variations and network connections, i.e., DNB, resulting in a distinct increase in the DNFE score, which indicates the upcoming tipping point or critical state.

To verify the robustness of the method, we compared the DNFE method with the sJSD method^14^ under different noise strengths in Fig 2D. As the noise intensity increases, the DNFE method outperforms the sJSD method in providing early-warning signals of critical transition stably, demonstrating that the DNFE method is significantly more effective and robust.

Furthermore, a 100-node and a 1000-node genetic regulatory network described by stochastic differential equations were used to generate two simulation datasets to identify the early-warning signals when the system approaches a tipping point. Fig 2E and F demonstrates the dynamical change of DNFE score for the 100-node network and the 1000-node network, respectively. There is an sudden increase of DNFE score at the parameter value *p* = 0, which suggests the imminent tipping point or critical state at a bifurcation point (*p* = 0). Clearly, when the system is far away from the tipping point, the DNFE scores of all nodes are steady and at a low level; when the system approaches the tipping point *p* = 0, there are significant increase of DNFE score around *p* = 0, i.e., the bifurcation point. So DNFE still maintains robust when it was tested by 1000-node networks, even much more network nodes. And when the algorithm was applied to the simulation dataset generated by 1000-node network, the result was calculated quickly which illustrated the application of the algorithm for large-scale or large quantities of data. The simulation and calculation details are presented in Supplementary Material: A.

### Identifying cell fate commitment during the cellular differentiation process

To demonstrate the computational efficiency of the proposed algorithm on single-cell datasets, we utilized the DNFE method on three scRNA-seq datasets of cell differentiation: hESC-to-DEC data, mESC-to-MP data, and MEF-to-Neurons data. The applications of DNFE method in hESC-to-DEC data is illustrated in the main text, and the results of mESC-to-MP data and MEF-to-Neurons data are provided in the Supplementary Material: B.

### Critical states for hESC-to-DEC

As depicted in Fig 3A, two critical states are identified with a sudden increase in the DNFE score, specifically, the first critical state at 12h and the second critical state at 36h, indicating two distinct differentiation processes post critical transitions, namely, ES-to-ME and ME-to-DE. The top 5% of genes with the highest DNFE score are DNBs, which are particularly sensitive to the critical state preceding disease deterioration. Fig 3B illustrates the landscape of global DNFE score, where the score significantly increases at 12h and 36h, suggesting the identified tipping points or critical states. Fig 3C displays the dynamic evolution of the Protein-Protein Interaction (PPI) directed network of DNFE signal biomarkers. It is evident that there is a major shift in DNFE score for the network post the critical point, implying that the network structure undergoes a significant change either from low to high, or vice versa. The nodes represent genes and directed edges represent the interactions between genes. Dynamic directed networks can provide information about gene interactions from multiple perspectives. Firstly, it can describe the temporal relationships among genes, which is essential for understanding the dynamic properties of interaction networks as well as the regulatory mechanisms. Secondly, dynamic directed networks can show the directionality of gene interactions, which is crucial for understanding the topology and signal transmission path of protein interaction networks. Moreover, as shown in Fig 3D and E, using the gene expressions of DNB genes cannot accurately distinguish the critical state from other states, but by using DNFE scores of those DNB genes, we can detect the critical state as shown in Figs 3A.

**Figure 3.**
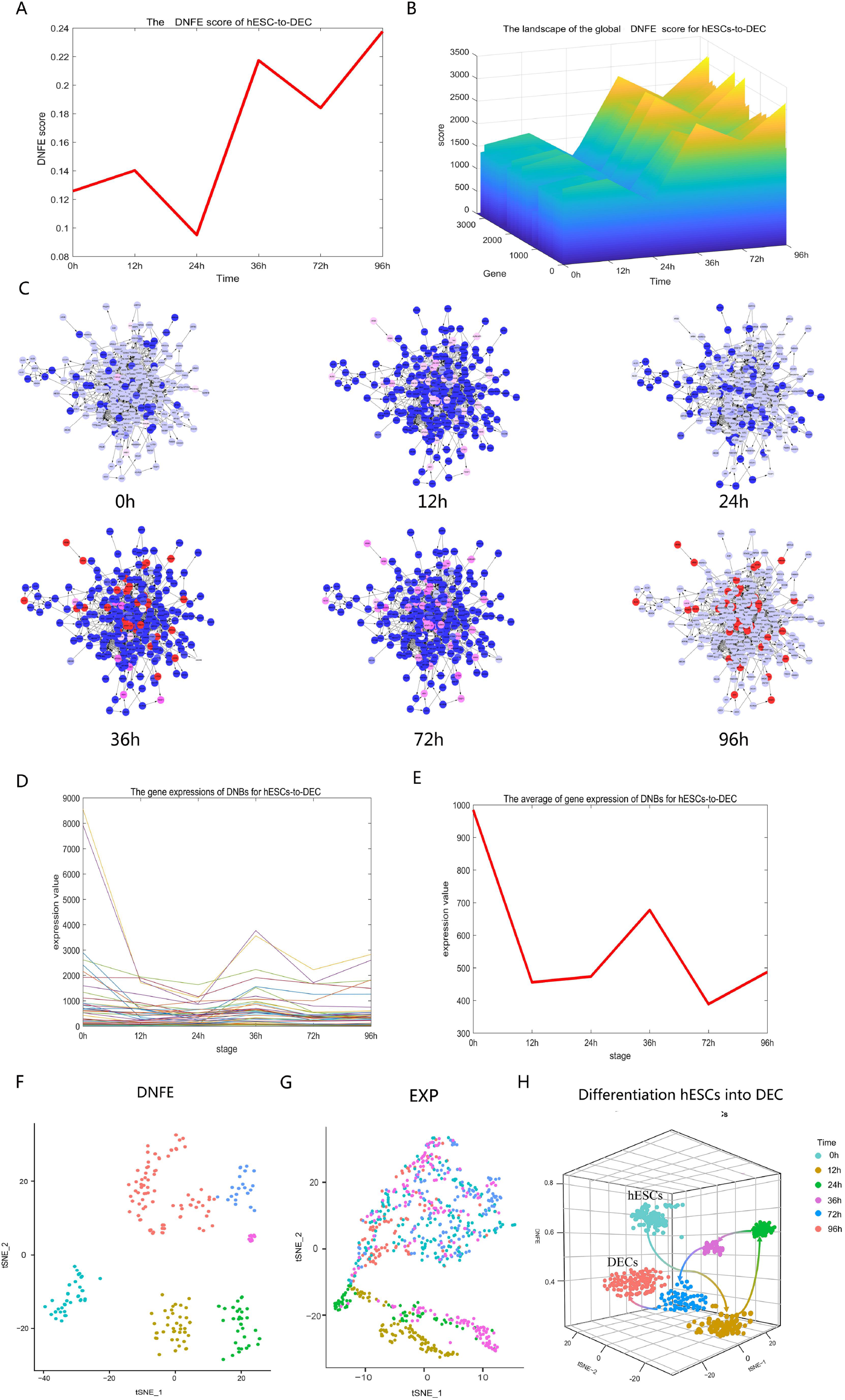
(A)DNFE scores at 12 and 36 h of hESCs’ differentiation are higher than those at other time points, which indicates that 12 and 36 h were the tipping points during the differentiation of hESCs. (B)The landscape of the global DNFE score for hESC-to-DEC. The score in DNBs arrives peak at 12h and 36h, indicating the two critical states. (C) The dynamic evolution of the gene directed regulatory network constructed by signaling genes for hESC-to-DEC data. The nodes represent genes and directed edges represent the interactions between genes. (D) The gene expression of DNBs for hESCs into DEC. (E) The average of gene expression of DNBs for hESCs into DEC. It is clear that the gene expression of DNBs cannot differentiate the critical state from other states, but DNFE method can detect the critical state. (F-H) Nodes in different colors represent cells from different time points. Clearly, DNFE distinguishes the temporal cell state better than EXP. The differentiation trajectories can be accurately predicted by DNFE scores.

The clustering analyses are shown in Fig 3F-3G, the clustering analysis based on DNFE can distinguish the state of cells at different time points while the gene expression fails. Furthermore, to further validate the DNFE performance, the pseudo-trajectory analysis was performed on the hESC-to-DEC data. Based on the temporal cell clustering by DNFE, the three-dimensional representations of cell-lineage trajectories are shown in Fig 3F, the z-axis represents DNFE potency estimation, while the x and y axes correspond to the t-SNE components. Fig 3H illustrates the developmental trajectories of cell differentiation from hESCs to DECs. The differentiation toward DECs appears after 36 h, which coheres with the experimental results^25^. These results demonstrate that the DNFE-based potency estimation can track the dynamic changes in cell potency, as well as the specific time point at which the cell fate commitment or the differentiation into distinct cell types occurs.

### Transcription factors in endodermal differentiation of hESCs

The top 5% of genes with the highest DNFE score are dynamic network biomarkers, where the genes that without differential expressions but are sensitive to DNFE score are known as ‘dark genes’. As expected, the DNB genes at 12 h showed higher scores than those of the two neighboring time points (0 and 24 h) in Fig 4A-4B, which suggests the two critical states, similarly to at 36 h in Fig 4C-4D.

**Figure 4.**
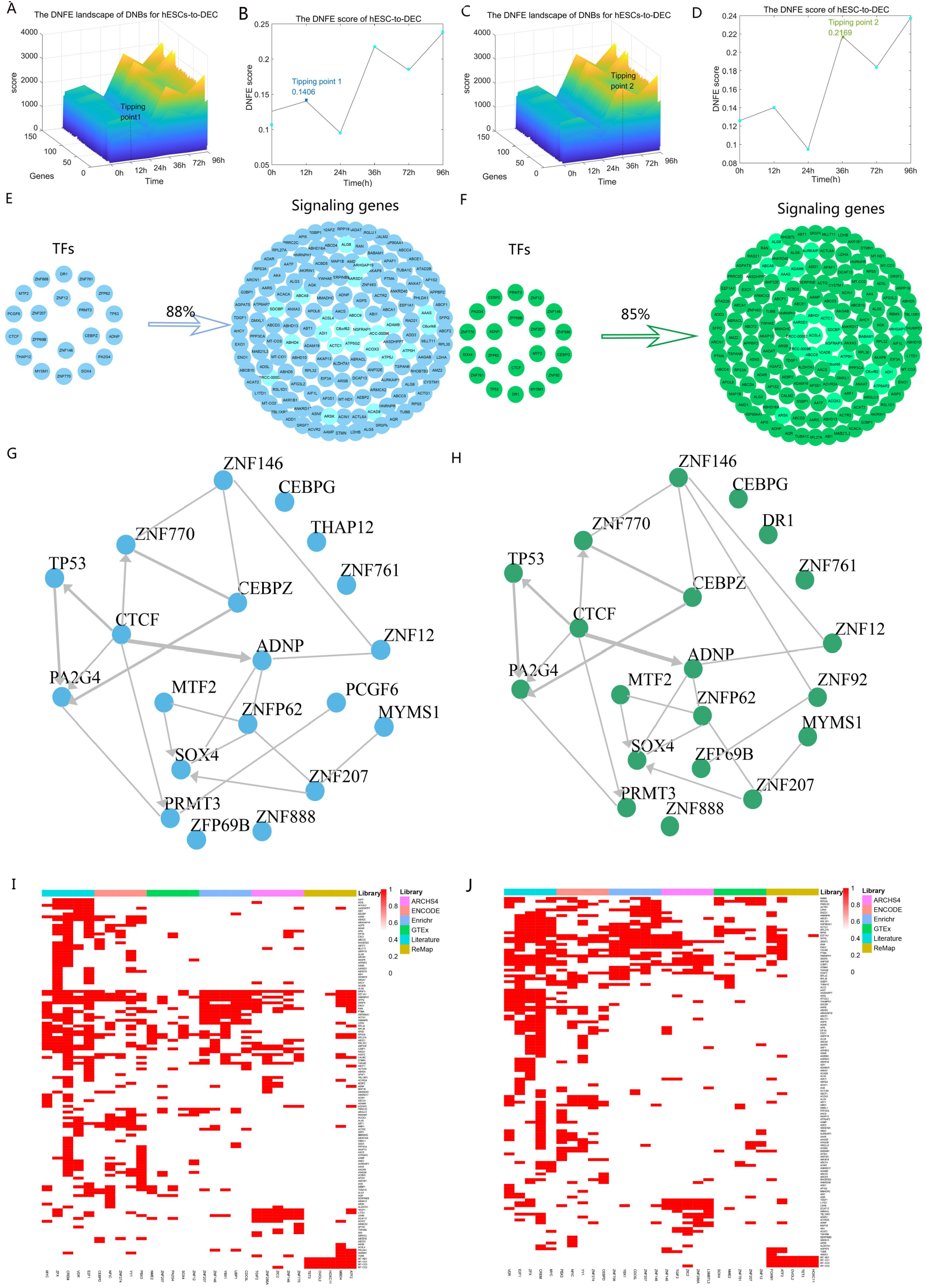
The tipping points of hESCs’ differentiation process revealed by the DNFE method. (A) Behavior of DNB genes at 12 h in the hESCs’ differentiation process. (B) DNFE scores at 12 h in the hESCs’ differentiation process. (C) Behavior of DNB genes at 36 h in the hESCs’ differentiation process. (D) DNFE scores at 36 h in the hESCs’ differentiation process. (E) Twenty hub upstream transcription factors could regulate 88% of DNB genes that were identified at 12 h (the first tipping point). (F) Twenty hub upstream transcription factors could regulate 85% of DNB genes at 36 h (the second tipping point). (G-H)The results of the regulation between the top 20 transcription factors at the first tipping point. (H) The results of the regulation between the top 20 transcription factors at the second tipping point. (I) The result with the top 5 TFs returned by each library on the columns and query genes on the rows is populated according to whether the query gene appears within the target gene set of the library TF at the first tipping point. (J) The result with the top 5 TFs returned by each library on the columns and query genes on the rows is populated according to whether the query gene appears within the target gene set of the library TF at the second tipping point.

Then, we set out to identify the potential upstream transcriptional regulators of the DNB genes of the two tipping points. Transcription factors (TFs) were focused because they are key players that define the cell identity and drive cell-fate transitions. We predicted the TFs on the chEA3 website, and identified twenty TFs for the first tipping point (12 h) and twenty TFs for the second tipping point (36 h) based on the chosen top 150 DNBs, respectively. These TFs of two groups could regulate 88% and 85% of the DNB in each tipping point, respectively (Fig 4E-4F). Among these DNB factors, ZNF146 is associated with poor cancer prognosis and plays a mitigating effect in their role as pro-proliferative and growth-sustaining transcriptional engines in cells^26^. CEBPZ is the key transcription factors that regulate various aspects of cellular differentiation and function in a variety of tissues^27^. PRMT3 overexpression promoted cell proliferation, migration, and invasion. Moreover, PRMT3 stabilized C-MYC and the pro-proliferation function of PRMT3 is dependent on C-MYC^28^. In hESCs, ZNF207 partners with master pluripotency TFs to govern self-renewal and pluripotency while simultaneously controlling commitment of cells towards ectoderm through direct regulation of neuronal TFs, which plays a important role during differentiation^29^. Clinically, these findings might provide a novel therapeutical treatment strategy for disease. As shown in Fig 4G, local network shows the results of the mutual regulation between the predicted top 20 transcription factors in the regulatory network, at the first tipping point, similarly to the second tipping point in Fig 4H. Especially, in the predicted top 20 transcription factors, we found ZNF888 which is key players that define the cell identity and drive cell-fate transitions, thus providing a theoretical evidence to understand the mechanisms of stem cell pluripotency and ESCs’ differentiation. However, mechanism of ZNF888 is ignored by traditional methods and may be a potential biomarkers, further experimental validation is required. As shown in Fig 4I-4J, heatmap demonstrates the result whether the gene appears within the target gene set of the library TF. The top five TFs are included in each library, and each column represent each TFs. SRSF3 and HNRNPH1 appeared most frequently in the target gene set of the library TF at the two tipping points. SRSF3 acted a role in termination of transcription and not in cleavage, maybe by interacting with the RNA downstream of the cleavage site^30^. In vivo and in vitro experiments showed that knockdown of HNRNPH1 inhibited cell proliferation and promoted cell apoptosis in CML cells^31^.

In summary, using the DNFE method, we identified two tipping points during the endodermal differentiation of hESCs and predicted twenty TFs that may play key regulatory roles in the cell-fate determination in this process.

### Potential biological functions of DNBs and nondifferential ‘dark genes’

To further explore the dark matter^32^ in the regulators of cancer-related pathways, we compared the DNB genes of disease with the differentially expressed genes. We found that there are some genes in the DNBs that are not differentially expressed at the molecular level but have a high DNFE score at the network level, which are known as ‘dark genes’.

To identify important pathways to characterize the potential biological mechanisms of ‘dark genes’ in hESC-to-DEC, we next conducted gene set enrichment analysis (GSEA). Detailed hallmark pathways enrichment analysis information was shown in Supplementary Table S1. As shown in Fig 5A, the crucial pathways included cAMP signaling pathway, Spliceosome, Pathogenic Escherichia coli infectic. In addition, Kyoto Encyclopedia of Genes and Genomes (KEGG) pathway enrichment analysis was performed to reveal the potential biological functions of the ‘dark genes’ and their 1st-order differentially expressed genes (DEGs) which are differentially expressed at the molecular level in the DNB genes. The enrichment results showed that these ‘dark genes’ and their DEG neighbors were mainly enriched in the JAK-STAT signaling pathway, cell cycle, pathways in cancer, PI3K-Akt signaling pathway, and other cancer related signaling pathways (Fig 5B and C). For instance, Antagonizing JAK-STAT signaling can obstruct the transformation of preneoplastic lesions into malignant tumors^33^. The PI3K-Akt signaling pathway is responsible for tumor cell differentiation, proliferation, and apoptosis^34^. Deregulation of the cell cycle leads to abnormal cell proliferation in cancer^35^. “Pathways in cancer” entails multiple signaling pathways and is closely related to cancer^36^. ysregulation of the cell cycle is an important malignant feature of cancer, and it may be directly related to cancer progression and metastasis^37^.

**Figure 5.**
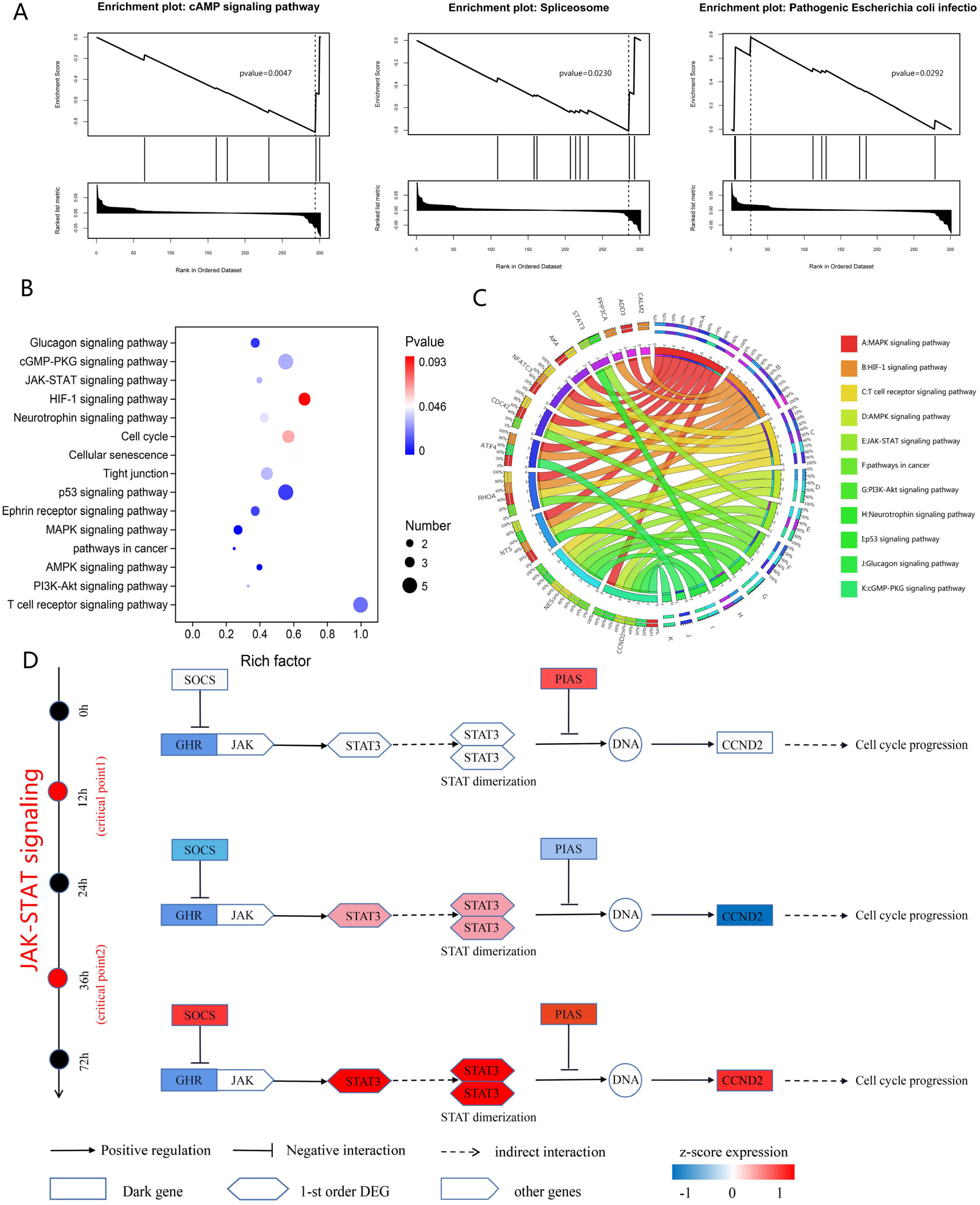
Regulatory mechanisms of embryo development revealed by the ‘dark genes’. (A) GSEA enrichment analysis of ‘dark genes’. The enriched pathways included cAMP signaling pathway, Spliceosome, Pathogenic Escherichia coli infectic. (B) KEGG pathway enrichment analysis for the ‘dark genes’ in hESC-to-DEC process. (C) DNBs are involved in important biological processes and KEGG pathways for in hESC-to-DEC process. The left side of the outer ring represents DNBs detected by DNFE, and the right side of the outer ring represents various biological processes in which these genes are involved. In the inner ring, the color and width of links indicate diverse enrichment pathways and significant levels of gene functions, respectively. (D) The underlying signaling mechanisms revealed by ‘dark genes’ and their 1st-order DEG neighbors.

Our results show that the synergy of the ‘dark genes’ and their 1st-order neighboring DEGs regulate cell cycle progression around the critical point through the JAK-STAT pathway (Fig 5D). The DNB GHR (growth hormone receptor), as the most upstream signaling molecule, did not show a significant change in gene expression level and is presented as a ‘dark gene’ from the perspective of DNFE^38^. Suppressors of cytokine signaling (Socs) genes inhibit GHR activation of STAT3^39^. The 1st-order neighboring DEG STAT3 changes significantly before and after the critical point, which may cause dimerization of STAT3, which in turn enters the nucleus and directly regulates the expression of downstream genes^40^. Protein inhibitor of activated STATs (PIAS) proteins were identified as negative regulators of cytokine signalling that inhibit the activity of STAT-transcription factors^41^. Furthermore, our results showed that CCND2 gene expression changed significantly after the critical point, which may promote the progression of the cell cycle in cancer cells^42^. These phenomena are consistent with our hypothesis that cell cycle progression signals are transmitted sequentially throughout the cancer period through the JAK-STAT pathway, with the final effect being observed only after the critical point transition.

### Identifying the critical transition during tumor progression

To demonstrate the computational efficacy of our proposed algorithm, we utilized the DNFE method on two tumor datasets (KIRP, BLCA) from the TCGA database. The applications of DNFE method in KIRP is illustrated in the main text, and the results of BLCA is provided in the Supplementary Material: C.

### The detection of critical states for KIRP

Fig 6A reveals a sudden increase at stage II, indicating a critical state or a tipping point. This is followed by distant metastasis of cancer, leading to a significant deterioration in the patient’s tumor state. The top 5% of genes with the highest DNFE scores at the critical state were identified as DNBs. Fig 6B shows the critical state at the molecular level, and the DNFE score in DNBs increases significantly at stage II. Cancer-related mortality is primarily due to tumor deterioration, such as cancer metastasis, which involves the migration of carcinoma cells from a tumor to a distant site^43^. Therefore, detecting the critical state is crucial to prevent or prepare for impending deterioration, enabling timely and appropriate clinical interventions. Prognostic analysis was performed based on the clinical information of KIRP samples and the survival times of samples before and after the critical state (stage II). Fig 6C shows a significant difference in the survival curves of KIRP before and after stage II. Moreover, the survival times of the samples before the critical state are significantly longer than those after the critical state. The sudden deterioration of survival times in patients strongly suggests that stage II is the critical state of KIRP. The development of KIRP is a multistage process, with the risk of deterioration increasing from one stage to the next. In this study, we further explored the deterioration risk of KIRP by dividing the samples of each stage into two groups based on the median DNFE score of samples at different stages: the high-score samples and the low-score samples. We then compared the prognosis of the two groups of samples. Fig 6D shows that the red curve represents the survival probability of the high-score samples, while the light blue curve represents the survival probability of the low-score samples. From a statistical perspective, the p-values in KIRP prognosis analysis were 0.00084 (Stage I-II), 0.019 (Stage III), and 0.036 (Stage IV). The log-rank test p-values of prognosis analysis in the Kaplan-Meier plot are all less than 0.05, indicating that the prognosis analysis is statistically significant. Furthermore, the survival time of the high-score group at Stages I-II is higher than that of the low-score group. The survival time of the high-score group after the critical state is significantly lower than the low score group. The high-score samples after the critical state have higher risks of disease deterioration and poor prognosis. The prediction of deterioration risk can assist in early clinical treatment. Fig 6E shows the DNB molecular directed network regulated by genes, and the network structure of stage II is noticeably different from other stages. All these results confirm that stage II is the critical state. Furthermore, as shown in Fig 6F and 6G, we can distinguish the critical state from other states using DNFE scores of DNB genes (Fig 6A), while merely using the gene expressions of DNB genes fails.

**Figure 6.**
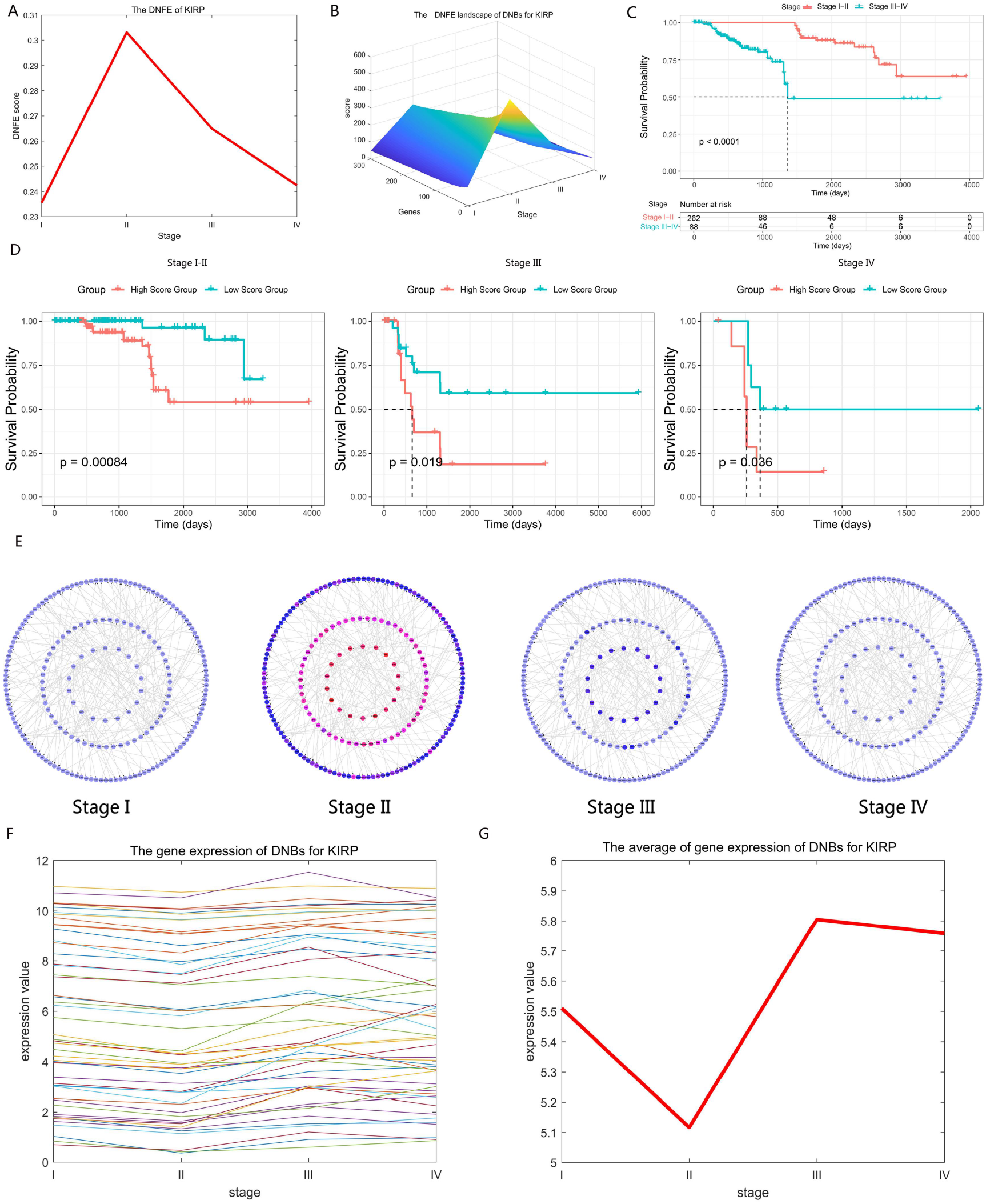
Identification of critical stage for KIRP. (A) The DNFE score curve of KIRP. The significant increase of DNFE score indicates the tipping point at stage II. (B) The DNFE landscape of DNBs for KIRP. The DNFE score increases significantly at stage II, indicating the critical state. (C) Comparison of survival curves for KIRP before and after the critical state/stage. The survival time before stage II is significantly longer than those after stage II. (D) Comparison of survival curves at different stages for KIRP between high-score samples and low-score samples. Red curves are the survival curves of high-score samples, and light blue curves are the survival curves of low-score samples. (E)The dynamic evolution of the DNB directed network structure for KIRP. The dynamic evolution of DNBs shows that the early-warning signals can be detected at stage II. (F) The gene expression of DNBs for KIRP. (G) The average of gene expression of DNBs for KIRP. It is clear that the gene expression of DNBs cannot distinguish the critical state from other states, but DNFE method can detect the critical state.

### The underlying signaling mechanisms revealed by DNBs and nondifferential ‘dark genes’

In addition, the enrichment analysis of DNBs validated the significance of DNBs in the progression of KIRP (Fig 7A). At the critical state, DNBs are enriched in some signaling pathways that are closely related to the PI3K-Akt signaling pathway, MAPK signaling pathway, pathways in cancer, and some other signaling pathways. Fig 7B illustrates the underlying mechanism revealed by the functional analysis on ‘dark genes’ and their 1st-order neighbors. It should be noted that there is a signal chain that responds to the ‘dark genes’ in the MAPK and PI3K/Akt signaling pathways, both of which are important for cell proliferation. In the MAPK signaling pathway, an identified ‘dark gene’, MAPK9, is the key gene of the c-Jun N-terminal kinase subclass pathway and may induce a variety of upstream signals to cause cell proliferation and differentiation^44^.

**Figure 7.**
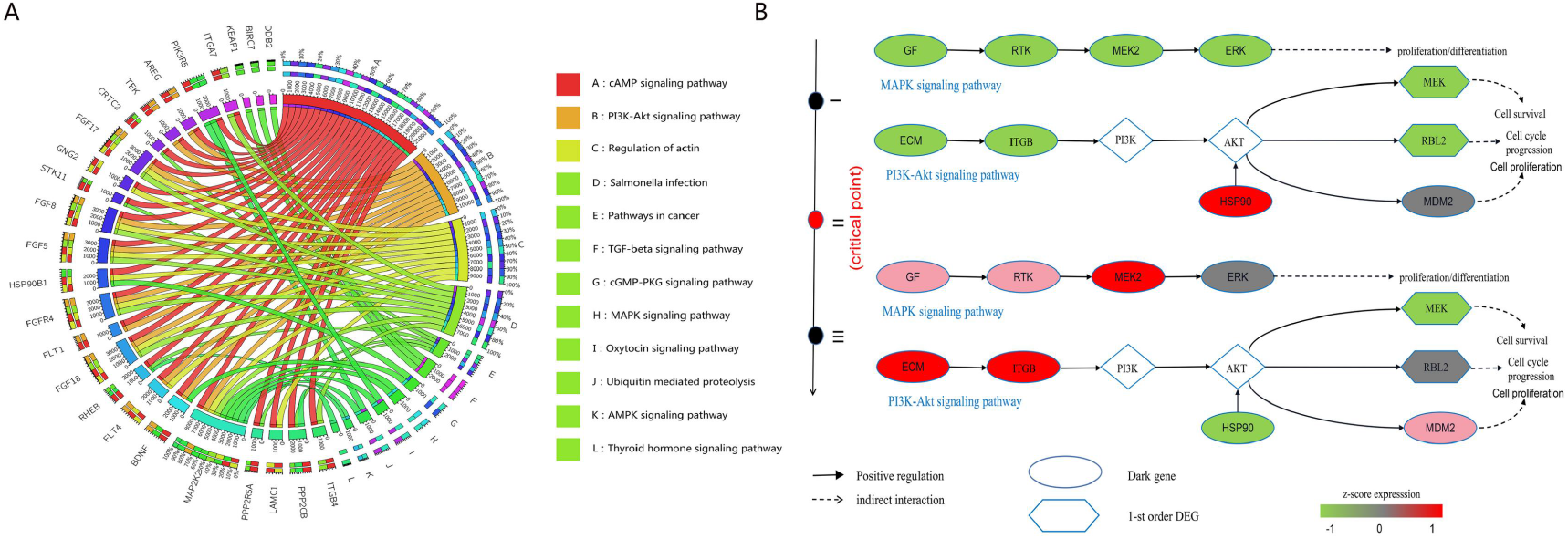
(A) DNBs are involved in important biological processes and KEGG pathways for KIRP. (B) Switching dynamics of DEGs before and after critical point induced by upstream ‘dark genes’. ‘Dark genes’ refer to the genes with nondifferential expression (P<0.05; t-test) but differential DNFE value (P<0.05).

In the PIK3/Akt signaling pathway, the upregulation of IGF1R, SOS1, and SOS2 will activate Ras and further activate RPS6KA3, which may cause the mitogenic effect^45^. The downstream signal response caused by ‘dark genes’ has a close relationship with the process of cell proliferation and differentiation. The accumulation of sustained expression of related genes in these pathways from stage I to stage III plays an important role in promoting proliferation and differentiation, which is consistent with the findings reported in the literature^46^ that the identified tipping point may be an important time point to guide the development of pluripotent stem cells to DE.

### Prognostic effects and functional roles of nondifferential ‘dark genes’

As shown in Fig 8, AKT1S1, NDRG1, VWF, ABCF3, TBC1D4, and PACSIN1 were found to be ‘dark genes’. When the critical state arrives, the DNFE score of ‘dark genes’ becomes more sensitive and shows obvious upward trends before the critical state to tumor deterioration compared to the gene expression data. Moreover, ‘dark genes’ also play an important role in cancer prognosis. We explored the effectiveness of ‘dark genes’ in prognosis by dividing all samples into two groups, i.e., a high-score group and a low-score group, based on gene expression and DNFE scores. As shown in Fig 8, for ‘dark gene’ VWF, there is an apparent increase in the DNFE score at the KIRP’s critical state, while its gene expression has no obvious fluctuations, indicating that the DNFE score can accurately capture the early warning signals of disease deterioration. In addition, the prognostic significance of the subgroup of ‘dark gene’ VWF based on the DNFE score and gene expression are 0.037 and 0.23, respectively, which means that the different subgroups obtained based on the DNFE score can stratify KIRP patients into subgroups with different survival times well from the viewpoint of statistics, while the gene expression fails, which indicates the effectiveness of the DNFE score in the early diagnosis of diseases. For other ‘dark genes’, we also obtained similar results.

**Figure 8.**
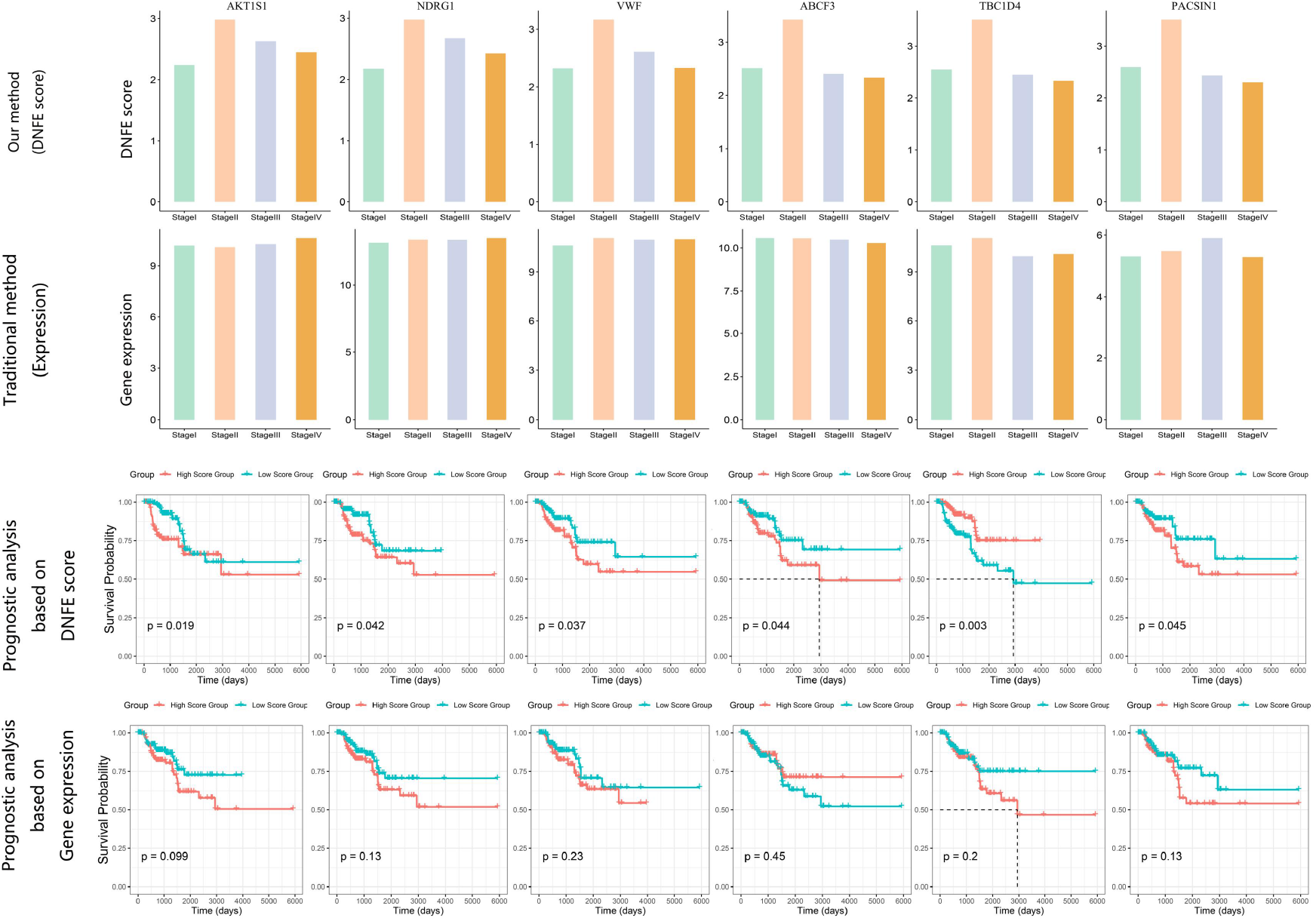
The comparison of prognosis analysis between gene expression and DNFE score of ‘dark genes’ for AKT1S1, NDRG1, VWF, ABCF3, TBC1D4 and PACSIN1, whose DNFE scores are more sensitive to the early warning signal of disease deterioration and can predict prognosis better than gene expression. The prognosis of these ‘dark genes’ shows that there are significant differences between the survival times of the two groups of samples, i.e., the high value group and the low value group, based on DNFE scores rather than gene expression.

Moreover, these ‘dark genes’ have significant functional roles in cancer processes. For example, downregulated expression of VWF in tumor tissues is associated with ERS model genes, which implies that it may play an important potential role in cancer progression^47^. EGFR-mediated signaling via PTPN11 (SHP2) and AKT1S1 (PRAS40) and an increase in anti-apoptotic signaling as a consequence of acquired cetuximab resistance provides a multitude of possibilities to identify and validate biomarkers, signaling pathways, and resistance mechanisms for clinically relevant improvement in cancer therapy^48^. Cell proliferation assays, colony formation assays, and cell migration assays show that overexpression of circRNA-TBC1D4 promotes NB cell migration, but not proliferation and colony-formation in vitro. These results suggest that circRNA-TBC1D4, often downregulated simultaneously in NB cell, may serve as biomarkers indicating an unfavorable NB type^49^.

## DISCUSSION

The identification of critical states in biological processes, such as the pre-deterioration stage of tumor diseases and cell fate commitment during embryonic development, holds significant importance. Early-warning signals for the critical state to disease states can provide optimal timing for prevention or preparation for catastrophic deterioration. However, the characterization of biological system dynamics and accurate detection of tipping points or critical states from datasets is challenging due to the similarity between pre-transition and critical states in terms of phenotype and mean gene expression. In this study, we propose a novel method based on a directed network to detect early-warning signals of biological processes, diverging from traditional methods that rely on differential expression information. Our results indicate that the proposed DNFE method can successfully detect the tipping point of five real datasets, identify DNBs, and reveal the underlying molecular mechanisms during disease progression. This method also enhances our understanding of the regulatory relationships between genes. We also discovered that ‘dark genes’ from three embryonic differentiation datasets and two cancer datasets play a significant role in crucial biological processes or pathways. TFs are pivotal in understanding cellular function, developmental biology, and disease mechanisms. In this study, we predicted TFs by identifying the ‘dark genes’ using web-based tools. However, this approach may not be entirely accurate, necessitating further experimental validation. Moreover, numerical simulations demonstrated the effectiveness and robustness of the DNFE method compared to existing methods, as well as the applicability to handle vast amounts of data or large-scale datasets.

In conclusion, we introduced an effective and robust computational method, DNFE, capable of detecting the tipping point of bulk and single-cell data and identifying corresponding DNBs. The DNFE method also shows promising potential for application in exploring potential molecular mechanisms of disease progression, discovering new network biomarkers, and ‘dark genes’.

## METHODS

### Data Progression and Functional Analysis

The DNFE method was applied to five real datasets, including three single-cell expression datasets and two bulk datasets.

The enrichment analysis of DNBs is based on the GeneOntology Consortium (http://geneontol-ogy.org), DAVID Bioinformatics Resources (https://david.ncifcrf.gov/) and Circos (http://www.circos.ca/). The transcription factors were predicted through CHEA3 (https://maayanlab.cloud/chea3/). Protein–Protein Interaction (PPI) networks were drawn using STRING (https://string-db.org/) and the client software Cytoscape (https://cytoscape.org/).

### Algorithm for disease prediction based on DNFE

The DNFE method is designed to detect the critical state or pre-disease state during the development of complex diseases and identify dynamic network biomarkers (DNBs). Given a group of reference/control samples representing the relatively normal/stable state, the following computational approach is designed to identify the critical state with only a specific sample. The schematic flowchart of the DNFE algorithm is demonstrated in Fig 1. The details of the procedure are described in the following subsections.

### [step1] Construct the directed network

Construct the WGCNA network based on reference samples and case samples. Based on WGCNA network and gene expression data, the global directed network can be constructed by a direction determination index, which is defined as follows.

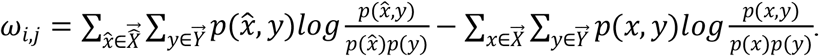

Vectors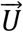 and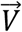 represent the expression profiles of genes *g*_*i*_ and *g*_*i*_ in all samples, respectively, 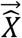 and 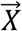 are defined as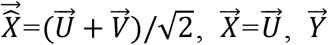 is the phenotype representing the binary vector for each sample (0-1). 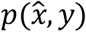 represents the joint probability density function (pdf) of 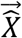 and 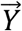, and *p*(*x,y*) represents the pdf of 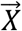 and 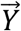, represent the edge pdf of 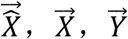, respectively. The positive determination value indicates that the integration of the gene *g*_*j*_ is an improvement of the gene *g*_*j*_ mutual information (MI), that is, in the directional network, there is a directed edge (*g*_*j*_, *g*_*j*_) from *g*_*i*_ to *g*_*j*_. Specifically, if *w*_*ij*_ is greater than zero, there is a directed edge (*g*_*i*_, *g*_*j*_) from gene *g*_*i*_, to *g*_*j*_; otherwise, there does not exist a directed edge (*g*_*i*_, *g*_*j*_). By this way, we construct the global directed network *N*^*global*^, where each directed edge (*g*_*i*_, *g*_*j*_) from gene *g*_*i*_ to *g*_*j*_ is decided by direction determination index *ω*_*ij*_. The theoretical background in network construction can be seen in Supplementary material: A.

### [step2] Extract gene module from global network

Each local gene module M^k^ = 1, …,m is extracted from global network. And, only first-order and second-order neighbors are considered in each gene module. Local gene module Mk centers on gene *g*^*k*^(*K* = 1,2, …, *m*) and *m*^*k*^ first-order out-degree neighbors 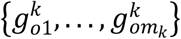, *n*^*k*^ first-order in-degree neighbors 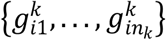

### [step3] Determine the local directed network

There are *p*_*j*_ second-order out-degree neighbors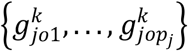, *q*_*j*_ second-order in-degree neighbors 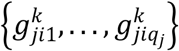 for each first-order neighborhood gene 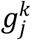. The cross-over strength of each first-order neighborhood gene 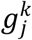 was calculated as follows,

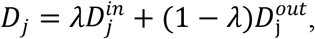

where λ is a constant, 0 < λ < 1, and the value can be determined on a case-by-case basis. In this paper, all algorithms are conducted under the principle of similarity weight, that is, the greater the weight of the edge, the smaller the distance between two points, and the closer the relationship. Therefore, the in-intensity of the node can reflect the importance of the node. 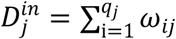 is the in-strength of the first-order neighborhood genes, and 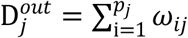 is the out-strength of the first-order neighborhood genes. *w*_*ij*_ represents the weight of the directed edge, (*g*_*i*_, *g*_*j*_) which is defined as

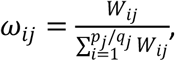

Where

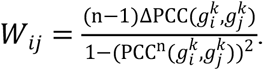

In this way, the crossover intensity of each first-order neighborhood gene 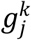 was ranked in descending order, and the top *l* first-order neighborhood genes were selected with the central gene *gk* to form a local central network *N*^*k*^(k = 1,2, …, *m*).

### [step4] Calculate a local DNFE score for each local directed network

We can obtain the local directed network *N*^*k*^(k = 1,2, …, *m*)., and each local directed network *N*^*k*^ is centered at gene *g*^*k*^, which has *a*^*k*^ first-order out-degree neighbors 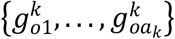 and first-order in-degree neighbors 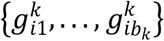, and *a*_*k*_+ + *b*_*k*_= *l*.

For the local directed network (with *a*_*k*_ first-order out-degree neighbors and first-order in-degree neighbors) centered on gene *g*^*k*^, its corresponding local DNFE score at t = T is defined below.

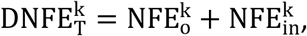

where 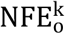 and 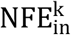 are defined as follows

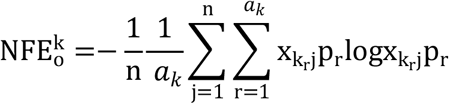

With

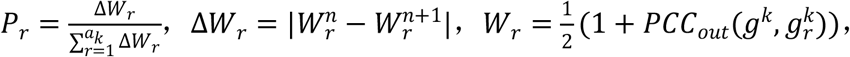

And

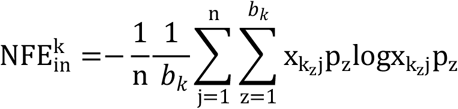

With

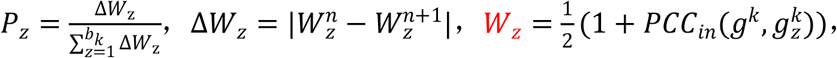

where *W*^*n*^ and ^1^ are the transformed Pearson correlation coefficients (PCCs) of gene expression between the center gene *g*^*k*^ and its first-order neighbor 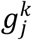 based on *n* reference samples and *n* 1 mixed samples (consisting of reference samples and one specific sample), respectively. and and one specific sample), respectively. 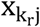 and 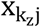are the expression data of gene are the expression data of gene 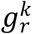 and 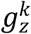 in sample *j*.

### [Step 5] Calculate the DNFE score of the global directed network

At time point t = T, the DNFE score for the perturbed sample is defined as follows,

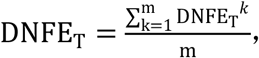

The DNFE score at each time point reflects the global perturbation caused by case samples. We repeat previous steps to calculate other score at different time t = T. Specifically, if score abruptly increase, T can be considered as an tipping point of critical transition during the system. We select the top 5% genes with the highest score at critical state as DNBs. The DNFE method mainly characterizes the network fluctuations in the system, which is the core of quantifying the criticality or tipping point of the network, thus providing a robust and reliable early warning signal for the critical state.

## Data availability

mESCs-to-MPs data data (ID:GSE79578), hESC-to-DEC data (ID:GSE75748), and MEF-to-Neurons data (ID:GSE67310) are accessible from the NCBI Gene Expression Omnibus (GEO) database (http://www.ncbi.nlm.nih.gov/geo). Kidney renal papillary cell carcinoma (KIRP) and bladder urothelial carcinoma (BLCA) are accessible from the cancer genome atlas (TCGA) database (http://cancergenome.nih.gov).

## Code availability

And the source code and a detailed usage guide of DNFE are freely available on GitHub now(https://github.com/pxqluck/DNFE).

## Author contributions

Xueqing Peng: Conceptualization, Methodology, Formal analysis, Validation, Visualization, Writing - original draft, Writing - review & editing. Peiluan Li: Conceptualization, Methodology, Writing - original draft, Writing - review & editing, Funding acquisition, Supervision. Luonan Chen: Conceptualization, Writing - review & editing, Funding acquisition, Supervision. All authors read and approved the final manuscript.

## Acknowledgements

This work was supported by National Key R&D Program of China (No. 2022YFA1004800), Strategic Priority Research Program of the Chinese Academy of Sciences (No. XDB38040400), National Natural Science Foundation of China (Nos. 12131020, 31930022, T2350003, T2341007), Special Fund for Science and Technology Innovation Strategy of Guangdong Province (Nos. 2021B0909050004, 2021B0909060002), Key-Area Research and Development Program of Guangdong Province (No. 2021B0909060002), JST Moonshot R&D (No. JPMJMS2021), and Major projects of Henan Province(NO.231100220100).

## REFERENCES

1. L.N. Chen, R. Liu, Z. Liu, et al., Detecting early-warning signals for sudden deterioration of complex diseases by dynamical network biomarkers, Sci. Rep. 2 (2012) 342.

2. Saelens W et al. (A comparison of single-cell trajectory inference methods. Nat Biotechnol 2019; 37:547–554.

3. Yuan Y, Bar-Joseph Z. Deep learning for inferring gene relationships from single-cell expression data. Proc Natl Acad Sci U S A 2019;116:27151–27158.

4. Abdelaal T, Michielsen L, Cats D et al. A comparison of automatic cell identification methods for single-cell RNA sequencing data. Genome Biol 2019;20:194.

5. Zhong, J. et al. (2021) SGE: predicting cell fate commitment during early embryonic development by single-cell graph entropy. Genomics Proteomics Bioinform., 19, 461–474.

6. Guo, W.F. et al. (2021a) Performance assessment of sample-specific networkccontrol methods for bulk and single-cell biological data analysis. PLoS Comput. Biol., 17, e1008962.

7. Shi, J. et al. (2021) Dynamics-based data science in biology. Natl. Sci. Rev., 8, nwab029.

8. R. Liu, K. Aihara, L. Chen, Dynamical network biomarkers for identifying critical transitions and their driving networks of biologic processes, Quant. Biol. 1 (2013) 105–114.

9. Gao, Rong et al. “Detecting the critical states during disease development based on temporal network flow entropy.” Briefings in bioinformatics vol. 23,5 (2022): bbac164. doi:10.1093/bib/bbac164.

10. P. Chen, R. Liu, L.N. Chen, et al., Identifying critical differentiation state of MCF-7 cells for breast cancer by dynamical network biomarkers, Front. Genet. 6 (2015) 252.

11. Yan, Jinling et al. “Disease Prediction by Network Information Gain on a Single Sample Basis.” Fundamental Research (2023): n. pag.

12. Zhong, Jiayuan et al. “Identifying the critical state of complex biological systems by the directed-network rank score method.” Bioinformatics (Oxford, England) vol. 38,24 (2022): 5398–5405. doi:10.1093/bioinformatics/btac707.

13. Yan, Jinling et al. “Identifying Critical States of Complex Diseases by Single-Sample Jensen-Shannon Divergence.” Frontiers in oncology vol. 11 684781. 4 Jun. 2021, doi:10.3389/fonc.2021.684781.

14. Liu, Xiaoping et al. “Detection for disease tipping points by landscape dynamic network biomarkers.” National science review vol. 6,4 (2019): 775–785. doi:10.1093/nsr/nwy162.

15. P. Chen, Y.J. Li, X. Liu, et al., Detecting the tipping points in a three-state model of complex diseases by temporal differential networks, J. Transl. Med. 15 (1) (2017) 217.

16. R. Liu, J.Y. Zhong, X. Yu, et al., Identifying Critical State of Complex Diseases by Single-Sample-Based Hidden Markov Model, Front. Genet. 10 (2019) 285.

17. M. Foo, J. Kim, D.G. Bates, Modelling and control of gene regulatory networks for perturbation mitigation, IEEE/ACM Trans. Comput. Biol. Bioinform. 16 (2) (2019) 583–595.

18. R. Khanin, V. Vinciotti, V. Mersinias, et al., Statistical reconstruction of transcription factor activity using Michaelis-Menten kinetics, Biometrics 63 (3) (2007) 816–823.

19. M. Ronen, R. Rosenberg, B.I. Shraiman, et al., Assigning numbers to the arrows: parameterizing a gene regulation network by using accurate expression kinetics, Proc. Natl. Acad. Sci. USA 99 (16) (2002) 10555–10560.

20. C. Sueyoshi, T. Naka, Stability analysis for the cellular signaling systems composed of two phosphorylation-dephosphorylation cyclic reactions, Comput. Mol. Biosci. 7 (2017) 33–45.

21. L. Chen, R. Wang, C. Li, et al., Modeling Biomolecular Networks in Cells: Structures and Dynamics, Springer, New York, 2010.

22. L. Chen, R. Wang, X. Zhang, Biomolecular Networks: Methods and Applications in Systems Biology, John Wiley & Sons, Hoboken, New Jersey, 2009.

23. Ronen M, Rosenberg R, Alon U, et al. Assigning numbers to the arrows: parameterizing a gene regulation network by using accurate ex-pression kinetics. Proc Natl Acad Sci U S A 2002;99(16):10555–60.

24. Chen L, Wang R, Li C, et al. Modeling Biomolecular Networks in Cells: Structures and Dynamics. New York: Sprimger, 2010.

25. Chu L, Leng N, Zhang J, et al. Single-cell RNA-seq reveals novel regulators of human embryonic stem cell 467 differentiation to definitive endoderm. Genome Biol 2016;17(1):173. doi: 10.1186/s13059-016-1033-x

26. Milazzo, Giorgio et al. “Single-Cell Sequencing Identifies Master Regulators Affected by Panobinostat in Neuroblastoma Cells.” Genes vol. 13,12 2240. 29 Nov. 2022, doi:10.3390/genes13122240.

27. Xia Y, Qadota H, Wang ZH, Liu P, Liu X, Ye KX, Matheny CJ, Berglund K, Yu SP, Drake D, Bennett DA, Wang XC, Yankner BA, Benian GM, Ye K. Neuronal C/EBPβ/AEP pathway shortens life span via selective GABAnergic neuronal degeneration by FOXO repression. Sci Adv. 2022 Apr;8(13):eabj8658. doi: 10.1126/sciadv.abj8658. Epub 2022 Mar 30. PMID: 35353567; PMCID: PMC8967231.

28. Hu, Yongbo et al. “Arginine methyltransferase PRMT3 promote tumorigenesis through regulating c-MYC stabilization in colorectal cancer.” Gene vol. 791 (2021): 145718. doi:10.1016/j.gene.2021.145718.

29. Fang F, Xia N, Angulo B, Carey J, Cady Z, Durruthy-Durruthy J, Bennett T, Sebastiano V, Reijo Pera RA. A distinct isoform of ZNF207 controls self-renewal and pluripotency of human embryonic stem cells. Nat Commun. 2018 Oct 22;9(1):4384. doi: 10.1038/s41467-018-06908-5. PMID: 30349051; PMCID: PMC6197280.

30. Zhou, Zhixia et al. “Emerging Roles of SRSF3 as a Therapeutic Target for Cancer.” Frontiers in oncology vol. 10 577636. 25 Sep. 2020, doi:10.3389/fonc.2020.577636

31. Liu M, Yang L, Liu X, Nie Z, Zhang X, Lu Y, Pan Y, Wang X, Luo J. HNRNPH1 Is a Novel Regulator Of Cellular Proliferation and Disease Progression in Chronic Myeloid Leukemia. Front Oncol. 2021 Jul 6;11:682859. doi: 10.3389/fonc.2021.682859. PMID: 34295818; PMCID: PMC8290130.

32. J.M. Johnson, S. Edwards, D. Shoemaker, et al., Dark matter in the genome: evidence of widespread transcription detected by microarray tiling experiments, Trends Genet. 21 (2) (2005) 93–102.

33. Bose S, Banerjee S, Mondal A, Chakraborty U, Pumarol J, Croley CR, et al. Targeting the JAK/STAT Signaling Pathway Using Phytocompounds for Cancer Prevention and Therapy. Cells (Basel, Switzerland). 2020;9(6):1451.

34. Martini M, De Santis MC, Braccini L, Gulluni F, Hirsch E. PI3K/AKT signaling pathway and cancer: An updated review. Annals of medicine (Helsinki). 2014;46(6):372–83.

35. Nakayama KI, Nakayama K. Regulation of the cell cycle by SCF-type ubiquitin ligases. Seminars in cell & developmental biology. 2005;16(3):323–33.

36. Dai Y, Qiang W, Yu X, Cai S, Lin K, Xie L, et al. Guizhi Fuling Decoction inhibiting the PI3K and MAPK pathways in breast cancer cells revealed by HTS2 technology and systems pharmacology. Computational and structural biotechnology journal. 2020;18:1121–36.

37. Hanahan D. Hallmarks of cancer: new dimensions[J]. Cancer discovery, 2022, 12(1): 31–46.

38. Zhong J, Han C, Zhang X, Chen P, Liu R. scGET: predicting cell fate transition during early embryonic development by single-cell graph entropy. Genomics, proteomics & bioinformatics. 2021;19(3):461–74.

39. Schaefer, F et al. “Impaired JAK-STAT signal transduction contributes to growth hormone resistance in chronic uremia.” The Journal of clinical investigation vol. 108,3 (2001): 467–75. doi:10.1172/JCI11895

40. Darnell Jr J E, Kerr M, Stark G R. Jak-STAT pathways and transcriptional activation in response to IFNs and other extracellular signaling proteins[J]. Science, 1994, 264(5164): 1415–1421.

41. Schmidt, D, and S Müller. “PIAS/SUMO: new partners in transcriptional regulation.” Cellular and molecular life sciences : CMLS vol. 60, 12 (2003): 2561–74. doi:10.1007/s00018-003-3129-1

42. Meyerson M, Harlow E D. Identification of G1 kinase activity for cdk6, a novel cyclin D partner[J]. Molecular and cellular biology, 1994, 14(3): 2077–2086

43. Zhong, Jiayuan et al. “SPNE: sample-perturbed network entropy for revealing critical states of complex biological systems.” Briefings in bioinformatics vol. 24, 2 (2023): bbad028. doi:10.1093/bib/bbad028

44. Zhang W, Liu HT. MAPK signal pathways in the regulation of cell proliferation in mammalian cells. Cell Res 2002;12:9–18.

45. Richardson CJ, Schalm SS, Blenis J. PI3-kinase and TOR: PIK-TORing cell growth. Semin Cell Dev Biol 2004;15:147–59.

46. Chu LF, Leng N, Zhang J, Hou Z, Mamott D, Vereide DT, et al. Single-cell RNA-seq reveals novel regulators of human embryonic stem cell differentiation to definitive endoderm. Genome Biol 2016;17:173.

47. Sadler, Brooke et al. “von Willebrand factor antigen levels are associated with burden of rare nonsynonymous variants in the VWF gene.” Blood vol. 137, 23 (2021): 3277–3283. doi:10.1182/blood.2020009999.

48. Wang, Chao et al. “Long Noncoding RNA LINC01134 Promotes Hepatocellular Carcinoma Metastasis via Activating AKT1S1 and NF-κB Signaling.” Frontiers in cell and developmental biology vol. 8 429. 12 Jun. 2020, doi:10.3389/fcell.2020.00429.

49. Eickelschulte, Samaneh et al. “AKT/AMPK-mediated phosphorylation of TBC1D4 disrupts the interaction with insulin-regulated aminopeptidase.” The Journal of biological chemistry vol. 296 (2021): 100637. doi:10.1016/j.jbc.2021.100637.

