## Supplemental file S1 for "DNFE: Directed-network flow entropy for detecting the tipping points during biological processes"

### Theoretical Background

Dynamic Network Biomarkers (DNBs) represent a collection of molecules, such as genes or proteins, or a molecular module that indicates the proximity of a tipping point or critical state, which is often followed by drastic deterioration connected with complex diseases[1]. As a biological or physiological system transitions from a stable normal state towards a critical state, a DNB module or a group of molecules (i.e., variables) emerge and meet the following three statistical conditions:

The standard deviation, representing deviation for each molecule within the module, drastically escalates;

The absolute values of the Pearson correlation coefficients among molecules within the module swiftly rise;

The absolute values of the Pearson correlation coefficients between molecules inside and outside of the modules markedly diminish.

These three conditions form the basis to detect the presence of DNBs or signals indicative of a pre-transition state. Moreover, these criteria are not arbitrary but embody the generic properties of DNB members whenever the system approaches a critical tipping point. These inherent properties have been effectively applied to numerous real diseases for detecting the tipping point or critical state.

The time-specific directed network at a time point T is constructed based on an information-theoretic scheme[2], which provides a direction determination index $\omega_{i,j}$ to evaluate the combined effect of gene combinations over a single gene from the perspective of mutual information (MI).

$$\omega_{ij}=\sum_{\hat{x}\in\vec{\hat{X}}} \sum_{y\in\vec{Y}} p(\hat{x},y)log\frac{p(\hat{x},y)}{p(\hat{x})p(y)}-\sum_{x\in\vec{X}} \sum_{y\in\vec{Y}} p(x,y)log\frac{p(x,y)}{p(x)p(y)}$$

Vectors $\vec{U}$ and $\vec{V}$ represent the expression profiles of genes $g_{i}$ and $g_{j}$ in all samples, respectively, $\vec{\hat{X}}$ and $\vec{X}$ are defined as $\vec{\hat{X}}$=$(\vec{U}+\vec{V})/\sqrt{2}$, $\vec{X}$=$\vec{U}$, $\vec{Y}$ is the phenotype representing the binary vector for each sample (0-1). $p(\hat{x},y)$ represents the probability density function (pdf) of $\vec{\hat{X}}$ and $\vec{Y}$, and $p(x,y)$ represents the pdf of$\vec{X}$and $\vec{Y}$, $p(\hat{x})$,$p(x),p(y)$represent the edge pdf of $\vec{\hat{X}}$，$\vec{X}$，$\vec{Y}$, respectively. The positive determination value indicates that the integration of the gene $g_{j}$ is an improvement of the gene $g_{i}$ mutual information (MI), that is, in the directional network, there is a directed edge ($g_{i},g_{j}$) from $g_{i}$ to $g_{j}$.

**Reference**

[1]Liu XP, Chang X, Leng SY, et al. Detection for disease tipping points by landscape dynamic network biomarkers. Natl Sci Rev 2019;6(4):775–85.
[2]Sun,D. et al. (2019) Discovering cooperative biomarkers for heterogeneous complex disease diagnoses. Brief. Bioinform., 20, 89–101.
